# Straightforward clustering of single-cell RNA-Seq data with t-SNE and DBSCAN

**DOI:** 10.1101/770388

**Authors:** Florian Wagner

## Abstract

Clustering of cells by cell type is arguably the most common and repetitive task encountered during the analysis of single-cell RNA-Seq data. However, as popular clustering methods operate largely independently of visualization techniques, the fine-tuning of clustering parameters can be unintuitive and time-consuming. Here, I propose *Galapagos*, a simple and effective clustering workflow based on t-SNE and DBSCAN that does not require a gene selection step. In practice, Galapagos only involves the fine-tuning of two parameters, which is straightforward, as clustering is performed directly on the t-SNE visualization results. Using peripheral blood mononuclear cells as a model tissue, I validate the effectiveness of Galapagos in different ways. First, I show that Galapagos generates clusters corresponding to all main cell types present. Then, I demonstrate that the t-SNE results are robust to parameter choices and initialization points. Next, I employ a simulation approach to show that clustering with Galapagos is accurate and robust to the high levels of technical noise present. Finally, to demonstrate Galapagos’ accuracy on real data, I compare clustering results to true cell type identities established using CITE-Seq data. In this context, I also provide an example of the primary limitation of Galapagos, namely the difficulty to resolve related cell types in cases where t-SNE fails to clearly separate the cells. Galapagos helps to make clustering scRNA-Seq data more intuitive and reproducible, and can be implemented in most programming languages with only a few lines of code.

## Introduction

Single-cell RNA-Seq is a breakthrough technology that provides a quantitative readout of the entire transcriptome of a cell, simultaneously for thousands of cells from a heterogeneous tissue or sample^1^. As the cell type identities of the captured cells are not known a priori, one typically has to perform clustering in order to assign a cell type identity to each cell. These clustering results then form the basis for a comparison of expression profiles between cell types or across biological conditions. The unique characteristics of single-cell RNA-Seq data, in particular the high intrinsic levels of technical noise^2^–^4^, have motivated the development of specialized scRNA-Seq clustering methods. A wide array of methods has been proposed, and a recent study has systematically compared twelve clustering methods on real and simulated data^5^.

Despite the large number of proposed methods, significant challenges have yet to be addressed: For example, clustering methods generally require a gene selection step, and different gene selection criteria can lead to very different clustering results^6^. However, little is known about how to choose the best gene selection criterion for a particular dataset. In addition, clustering methods often invoke a complex analysis workflow that depends on a large number of adjustable parameters, further increasing the range of possible results. Finally, clustering methods commonly operate independently of popular visualization algorithms such as t-SNE^7^ and UMAP^8^, thereby limiting the extent to which visualization results can guide the tuning of clustering parameters. In combination, these factors can make the process of clustering of scRNA-Seq data unintuitive and time-consuming, and can make the clustering results difficult to interpret and reproduce.

Here, I propose *Galapagos* (**G**enerally **a**pplicable **l**ow-complexity **ap**proach for the **ag**gregation **o**f **s**imilar cells), a simple clustering workflow for scRNA-Seq data designed to overcome the aforementioned challenges: First, it does not require gene selection, thus eliminating possible variation and bias from this step. Second, it consists of a simple series of steps with only four tunable parameters, whose individual roles are readily appreciated. Third, it performs clustering directly on visualization results, thus enabling a straightforward, visually guided tuning of clustering parameters. The Galapagos workflow can be easily implemented in most modern programming languages, and is intended to make clustering of scRNA-Seq data simpler, more transparent, and more reproducible.

## Results

### A straightforward approach to clustering single-cell RNA-Seq data

To design a straightforward approach for clustering single-cell RNA-Seq data, I relied on the observation that after median-normalization and variance-stabilization of the data^2,4,9^, *t-SNE^7^* appeared to reliably separate the main cell types in a dataset without requiring any type of gene selection. I furthermore observed that t-SNE tended to produce “islands” of cells with relatively even cell densities, and reasoned that this property could be effectively exploited by the density-based *DBSCAN* clustering algorithm^10^. This would enable the generation of clusters that correspond well to the visual groupings of cells that were readily apparent from t-SNE results. The resulting workflow, termed *Galapagos*, comprises five steps with a total of only four parameters (**Figure 1a**). In my experience, two of the parameters (*num_components* and *perplexity*) can be set to default values, and in most practical situations do not require any tuning at all. The remaining two parameters are those of the DBSCAN algorithm and can be tuned in a straightforward manner, using the t-SNE visualization result as a reference. A detailed description of the proposed workflow and tuning procedure can be found in the **Methods**.

To showcase the effectiveness of Galapagos, I applied the method to a scRNA-Seq dataset of human peripheral blood mononuclear cells (PBMCs), obtained from a healthy donor. PBMCs represent a useful model tissue for the development and benchmarking of scRNA-Seq clustering methods, as they contain a largely known composition of cell types with varying degrees of biological relatedness. Moreover, a relatively large number of PBMC scRNA-Seq datasets are publicly available, including datasets obtained with different scRNA-Seq platforms and protocols. The t-SNE plot produced by Galapagos for this particular dataset showed four main “cell islands” and a few small islands (**Figure 1b**, left panel). Additionally, some of the main islands appeared to be broken up into smaller islands in close proximity to each other (e.g., 4a and 4b in the figure). After tuning of the DBSCAN parameters (**Methods**), I obtained a clustering result that agreed well with the visual impression from the t-SNE result (**Supp. Figure 1a**). Based on a careful inspection of marker gene expression patterns, I annotated the clusters with ten different cell type identities, including three subtypes of T cells and two subtypes of B cells (**Figure 1b**, right panel). While the analysis of an isolated scRNA-Seq dataset did not allow for clustering accuracy to be quantified, an examination of the average expression levels of key marker genes showed that each cluster exhibited a clearly distinct expression profile, and that the cell assignments agree with known and previously reported expression patterns for each of the cell types (**Figure 1c and Supp Figure 1b)**. In summary, the application of Galapagos to PBMCs suggested that it was highly effective in distinguishing between the main cell types in the data.

**Figure 1:**
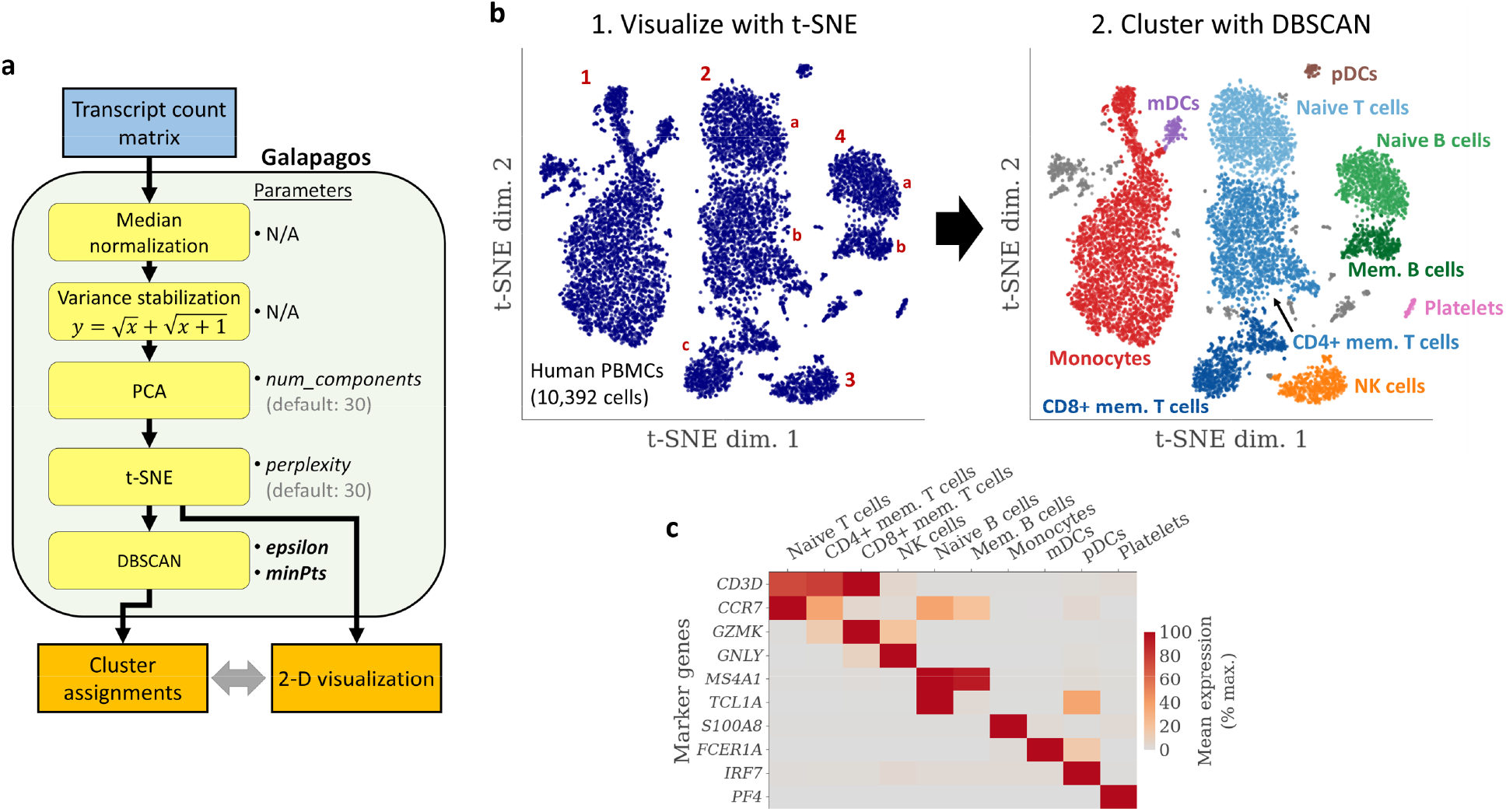
Galapagos workflow and application to human peripheral blood mononuclear cells (PBMCs). **a** Flow chart showing each step of the workflow and the associated parameters (if any). **b** t-SNE visualization (left) and DBSCAN clustering results (right) for a 10x Genomics PBMC dataset. Numbers indicate four main cell islands, and lower case letters indicate “sub-islands” in close proximity of each other. The workflow was run with default PCA and t-SNE parameters, and fine-tuned DBSCAN parameters (see main text). **c** Heatmap showing the mean normalized expression level of key marker genes in each of the clusters. For each gene, the expression levels are expressed in percent, relative to the maximum value across all clusters.

### Galapagos t-SNE results are robust to initialization points and parameter choices

Some researchers view t-SNE as a tool that is too unreliable or unstable to be used for clustering purposes. In the **Discussion**, I argue that while there is merit to the widespread skepticisms surrounding t-SNE results, the problem stems from the specific practice of how t-SNE analyses have come to be used in single-cell studies, not the t-SNE algorithm itself. One concern relating to the use of t-SNE is that the results are sensitive to initialization points and other random values generated during its execution. To provide evidence that t-SNE is robust to initialization points, even when analyzing complex datasets with more than 10,000 cells, I repeated the analysis presented in **Figure 1b** with different seeds for the random number generator. The results showed that while the exact location and the shape of individual cell islands changed, cluster assignments remained almost perfectly coherent across all analyses (**Figure 2**). Moreover, the clusters representing subtypes of T cells and B cells remained in close proximity, and the main characteristics of individual island were preserved. For example, some cells “protruded” from the monocyte island, which corresponded to a spectrum of CD14- and CD16-positive cells (not shown), and this protrusion was present in all analyses. The only noticeable difference was that the cluster of mDCs formed its own island in Replicate 2, rather than being in proximity to the monocyte island. This was likely an example of a borderline case where the differences between the two cell types were just large enough to prompt t-SNE to produce separate islands in some but not all t-SNE runs.

**Figure 2:**
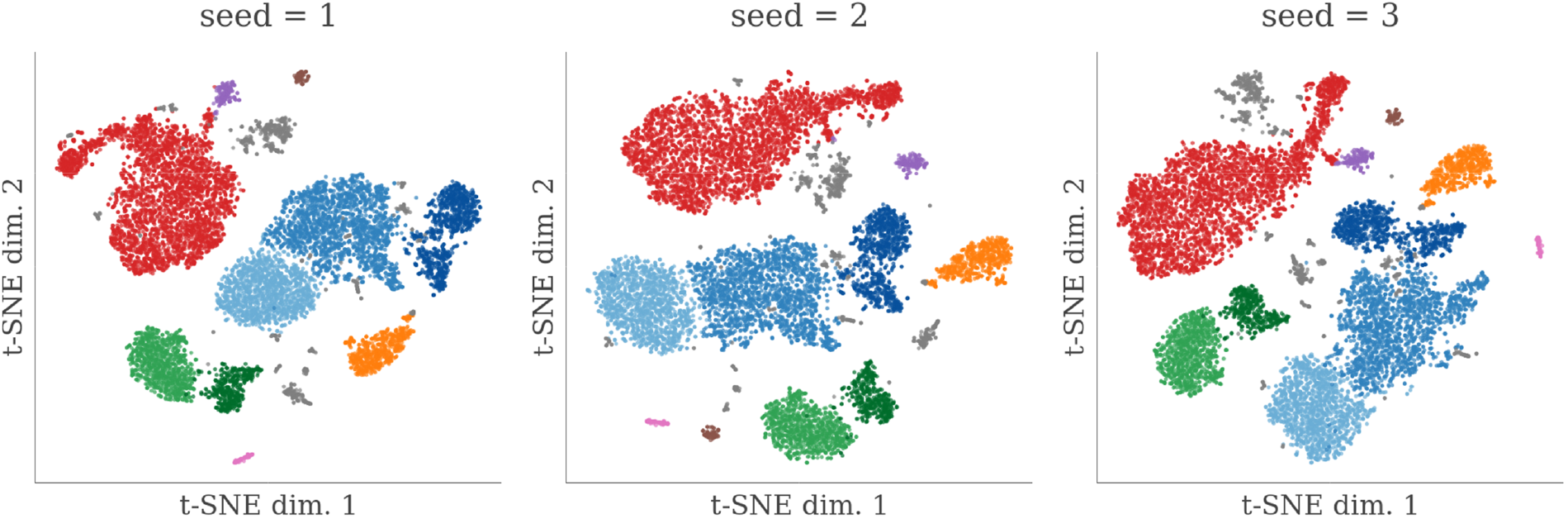
Robustness of Galapagos t-SNE results to initialization points. Shown are t-SNE results for the same PBMC dataset as in **Figure 1**, with three different initialization points (seed values). Overlaid onto t-SNE plot are the cluster assignments shown from **Figure 1b** (right). While different initialization points result in different geometries and spatial arrangements of the clusters, the cluster assignments are almost perfectly “coherent” across analyses.

Using a similar approach, I also repeated the analysis with different combinations of values for the *num_components* and *perplexity* parameters. The results showed that t-SNE results were highly robust to these parameters, again showing very little variation in the number, characteristic shape, or cell composition of the cell islands generated (**Supp. Figure 2**). In summary, t-SNE results were highly robust to both initialization points and parameter choices, thus justifying the use of t-SNE for clustering purposes.

### Galapagos accurately separates all major cell types in simulated scRNA-Seq data

The analyses presented above established that Galapagos recovered clusters corresponding to all main cell types present in PBMCs. However, as the true cell type identities of the cells in the dataset were not known, it was not possible to determine the clustering accuracy of the method. To overcome this limitation, I decided to adopt *Sim-ENHANCE*, a previously described approach for simulating scRNA-Seq datasets with realistic biological and technical characteristics^11^ (**Figure 3a**). First, I generated a “ground truth” dataset by applying the ENHANCE denoising algorithm^11^ to the real PBMC dataset. The application of Galapagos to the ground truth resulted in 10 clusters whose shapes resembled the results obtained for the real dataset (**Supp. Figure 3a**). As the simulation preserved a 1:1 relationship between cells in the real dataset and the ground truth, I was able to confirm that the ground truth recapitulated the cell type heterogeneity present in the real data by overlaying the original clustering results (**Supp. Figure 3b**). I then identified the cell type of each cluster, and treated the results as the “true” cell type identities (**Figure 3b**). An examination of marker gene expression patterns confirmed that the ground truth and true cell type identities closely resembled real PBMC data (**Figure 3c** and **Supp. Figure 3c**). Finally, to obtain realistic scRNA-Seq datasets based on the ground truth, I simulated technical noise, comprised of efficiency noise and sampling noise. The simulated datasets represented a set of technical replicates, as their technical noise was generated independently of each other. The introduction of technical noise tripled of the total variance contained in the data, resulting in levels that were comparable to the PBMC dataset that formed the basis for the simulation (**Figure 3d**). In summary, approximately one third the simulated datasets represented the biological cell type differences contained in the ground truth, and the other two thirds represented random technical noise.

**Figure 3:**
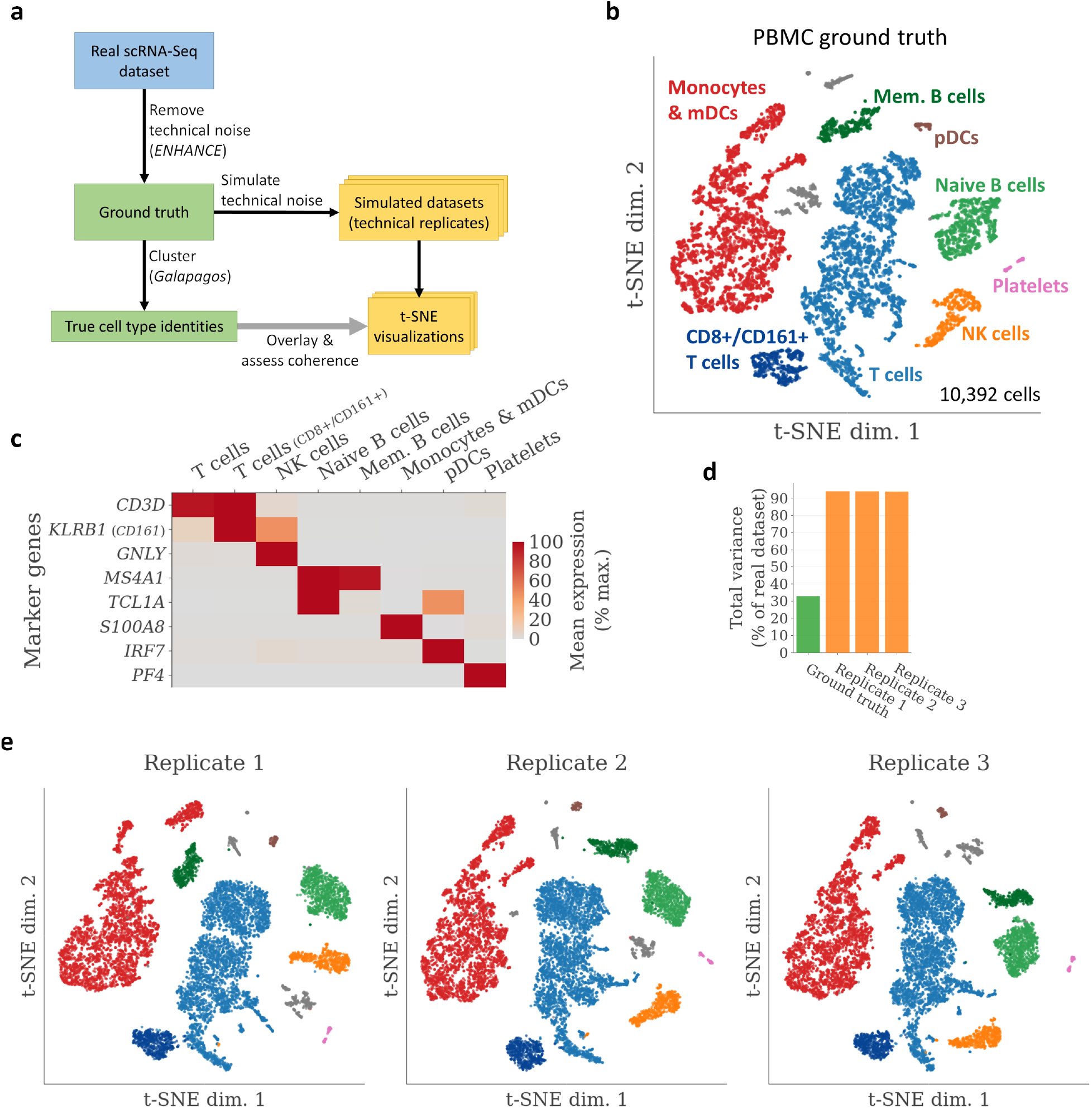
Examination of Galapagos clustering accuracy on simulated PBMC data. **a** Design of the simulation study. **b** Application of Galapagos to the ground truth. Shown are clustering results as well as manual cell type annotations. **c** Heatmap showing the relative expression levels of key marker genes, as in **Figure 2b**. **d** Comparison of the total amount of variance present in the simulated datasets. **e** Galapagos t-SNE visualizations of the three simulated datasets, overlaid with the true cell type assignments shown in (**b**).

I then applied the Galapagos workflow to the simulated datasets and overlaid the true cell type identities onto the t-SNE visualizations (**Figure 3e**). The results showed that Galapagos was indeed able to accurately recover cell islands that were highly consistent with the true cell type assignments, and that it did so reliably for all three replicates. A subset of cells from the monocyte & mDC cluster formed its own island in one of the replicates, again highlighting that the “breaking off” of sub-islands can be a somewhat stochastic event in t-SNE analyses. In summary, these analyses demonstrated that Galapagos was able to accurately separate all major cell types present in the simulated data, demonstrating its ability to overcome the high levels of technical noise present in scRNA-Seq data.

### Galapagos results closely agree with true cell type identities established using CITE-Seq data

After establishing that Galapagos could accurately identify clusters of cells in a simulation study, I intended to quantify the clustering accuracy of Galapagos on real data, based on an experimentally derived ground truth. To do so, I decided to take advantage of a CITE-Seq PBMC dataset that contained protein measurements for a panel of cell surface markers, in addition to the mRNA expression levels contained in regular scRNA-Seq data (**Figure 4a**). As the protein expression measurements were much less noisy than the mRNA data measurements, I was able to rely on simple thresholds for known marker genes in order to define T cells, NK, and monocytes (**Figure 4b**). This successive application of thresholds resembled the “gating” of cells typically used in analyzing flow cytometry data, and resulted in experimentally derived “true” cell type assignments that were independent of the mRNA expression data. I then applied Galapagos to the mRNA data and compared the clustering results with the true cell type assignments (**Figure 4c,d**). The results showed that T cells, NK cells, and monocytes were all identified with >95% precision and >90% recall, providing experimental evidence for Galapagos’ high clustering accuracy (**Figure 4e**).

### Galapagos is limited in its ability to separate cells belonging to closely related cell types

While the simulation study and the analysis of CITE-Seq data demonstrated Galapagos’ ability to accurately separate major cell types, it was less clear whether the method was able to clearly distinguish between closely related cell types or subtypes. To directly investigate this issue, I again used markers with protein expression measurements in the CITE-Seq PBMC dataset to distinguish between naïve and memory T cells, as well as CD4+ and CD8+ subsets of these T cell subtypes (**Figure 5a**). I then overlaid the cell type assignments on the original t-SNE visualization, and found that while the cells of each cell type appeared spatially coherent, the CD4+ and CD8+ naïve T cells were not separated from each other, thus making it all but impossible to distinguish them using the DBSCAN algorithm (**Figure 5b** and **Supp. Figure 4**). This provided a clear example of Galapagos’ main limitation, namely the difficulty to distinguish between cell types in cases where t-SNE fails to clearly separate the cell populations. This can be expected to occur whenever the expression differences between the cell types (here, CD4+ and CD8+ naïve T cells) are too small in relation to the level of noise affecting the expression measurements.

Re-applying Galapagos to only the T cells led to only marginally improved t-SNE results (**Figure 5c**). I also tried to replace t-SNE with the more recently developed UMAP algorithm, to test whether it would result in a better separation of CD4+ and CD8+ naïve T cells. However, the results showed a substantial overlap of these cell populations, indicating that UMAP performed worse than t-SNE in separating these subtypes of T cells. These analyses suggested that an entirely different clustering approach might be required in order to distinguish between closely related cell types. Based on previous work, an effective strategy in such situations could consist of computational denoising, followed by hierarchical clustering of highly variable genes^11^.

**Figure 4:**
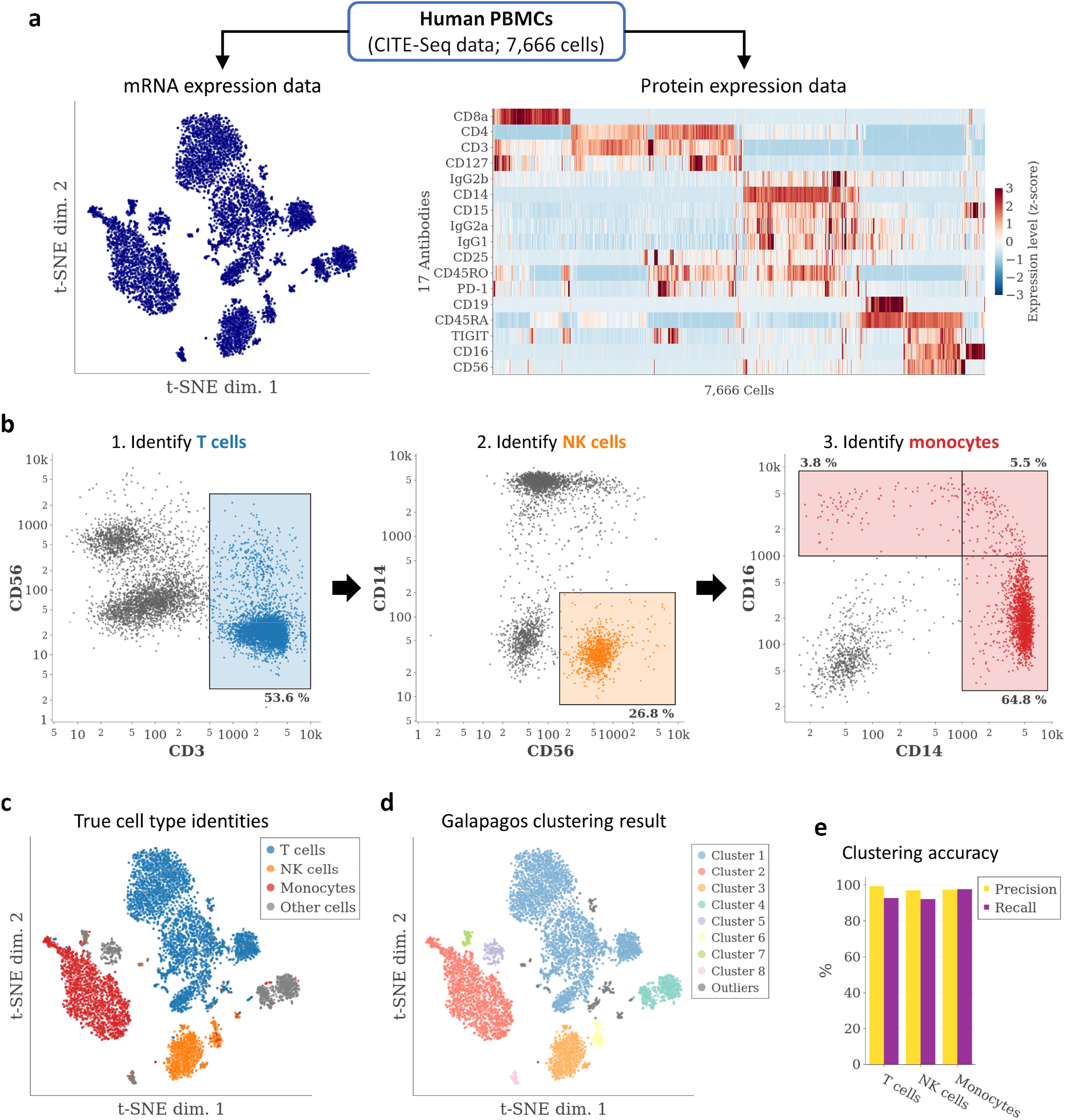
Quantification of clustering accuracy using an experimental ground truth. **a** t-SNE visualization of the mRNA expression data (left) and heatmap showing the protein expression data (right). Proteins and cells in the heatmap were ordered based on hierarchical clustering results. **b** Identification of T cells, NK cells, and monocytes based on the protein expression levels. The highlighted boxes show the “gates”, i.e., expression thresholds, that were applied sequentially to identify the indicated cell types. **c** t-SNE result from (**a**), overlaid with the cell type identities established in (**b**). **d** t-SNE result from (**a**), overlaid with the DBSCAN clustering result. **e** Precision and recall for each cell type, calculated by comparing the clustering results (**d**) with the true cell type identities (**c**).

**Figure 5:**
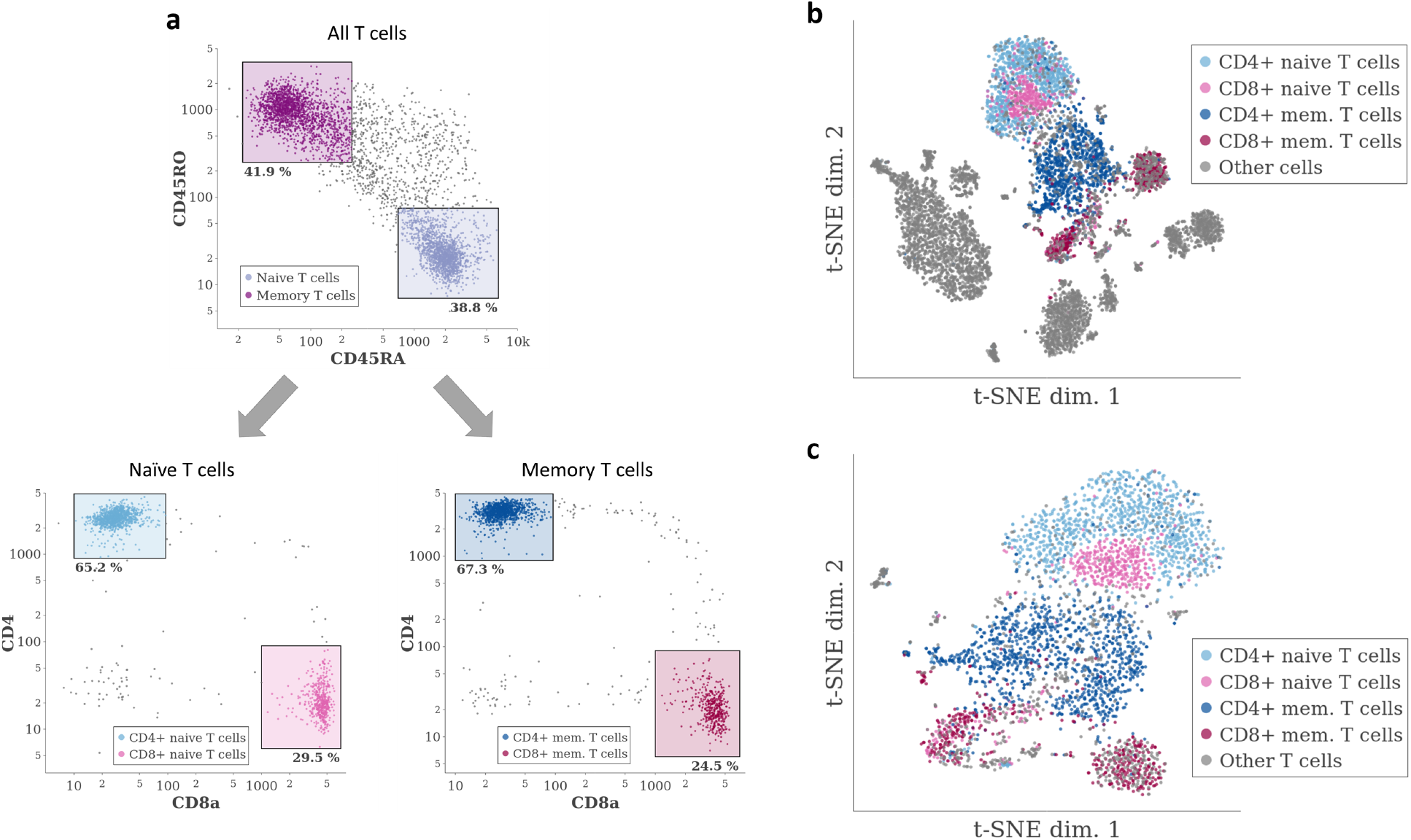
Galapagos’ resolution is limited by t-SNE’s ability to separate cells belonging to related cell types. **a** Identification of T cell subtype identities using protein expression measurements from the PBMC CITE-Seq dataset. **b** Original t-SNE result from **Figure 1b**, overlaid with the T cell subtype identities established in (**a**). **c** t-SNE result obtained after selecting only the cells identified as T cells in **Figure 1c**, overlaid with the same cell type identities as in (**b**).

## Discussion

I have described and validated *Galapagos*, a general-purpose method for clustering scRNA-Seq data that differs from currently popular clustering methods primarily by its simplicity, involving only a small number of analysis steps with few parameters, and its reliance on t-SNE visualization results. The main motivation for this work was the observation that despite the ubiquitous need for clustering scRNA-Seq data, obtaining satisfactory clustering results can still be a non-intuitive and time-consuming process. In addition, clustering workflows that operate independently from visualization techniques and involve many adjustable parameters produce results that are difficult to interpret and communicate. The Galapagos workflow was designed to be both simple and effective, thus making it easier to obtain satisfactory results. At the same time, the fact that clustering is performed directly on the t-SNE visualization makes it easier to communicate clustering results, and the limited number of analysis steps and parameter choices result in increased transparency and reproducibility. I believe that these features make Galapagos an attractive choice for visualizing and clustering scRNA-Seq data.

Despite its ubiquitous use as a method for visualization scRNA-Seq data, t-SNE has rarely been proposed for use in clustering workflows. In their systematic evaluation of clustering methods, Duò et al. included a method in which cells are clustered by applying the k-means algorithm to t-SNE results. However, k-means is not a suitable choice for clustering data after applying a non-linear dimensionality reduction method such as t-SNE that only preserves local, but not global distances^5^. It is sometimes believed that t-SNE results have merely “illustrative” character and are not sufficiently reliable to justify their use in clustering. Additionally, it is not uncommon for researchers to place much more trust in the t-SNE results they have generated themselves, than in those generated by others. I believe much of this skepticism is the result of the myriad different ways in which the scRNA-Seq data is normalized, filtered, and transformed, prior to the application of t-SNE. It can be difficult even for experienced researchers to predict how different approaches affect the visualization result, and it is therefore sensible to apply a high degree of skepticism toward any particular t-SNE plot. It is interesting to note that these attitudes parallel skepticisms about the reliance on p-values to decide whether a particular treatment or condition had a desired effect. As Leek and Peng have pointed out, p-values are generally only the “tip of the iceberg”, the result of a complex sequence of analysis steps that are rarely scrutinized^12^. t-SNE plots similarly represent the tip of the iceberg in the visualization of scRNA-Seq data. I have therefore adopted the view that the solution to the perceived lack of t-SNE reliability and robustness is not to avoid using t-SNE altogether, but rather to simplify and standardize the way in which the data are preprocessed. This greatly reduces the chance of results being biased, increases the transparency of the analysis, and hopefully allows researchers to place greater trust in t-SNE plots and clustering results that they have not generated themselves.

UMAP has recently gained popularity as an alternative to t-SNE for visualization single-cell data^8,13^. While a formal comparison of UMAP and t-SNE performance for clustering scRNA-Seq data was beyond the scope of this article, I currently do not believe that UMAP would be a useful alternative to t-SNE in the context of the Galapagos workflow. Specifically, it does not appear that UMAP visualizations exhibit an equally even cell density across cell islands, which is the key property that enables the effective use of DBSCAN. I would therefore anticipate that it is more difficult to effectively apply DBSCAN to UMAP results than to t-SNE results. Moreover, it is currently unclear whether UMAP provides a qualitatively better separation of cell types than t-SNE. Notably, the authors of UMAP have pointed out the similarities between the two methods, and described UMAP as belonging to the same class of “k-neighbour based graph learning algorithms” as t-SNE. A key advantage of UMAP over t-SNE appears to be its significantly faster runtimes on large datasets, however a recently developed variant of the t-SNE algorithm has been shown to have similarly fast runtimes in some cases^8,13,14^.

The present study has a few notable limitations. I did not directly compare the complexity, robustness or accuracy of Galapagos and previously proposed clustering methods. While comprehensive comparisons are laborious, given the large number of proposed methods^5^, it would be useful to compare Galapagos directly to some of the more popular approaches. Moreover, I did not present validations of Galapagos on tissues other than PBMCs, which would provide evidence for its effectiveness independently of the tissue analyzed. Preliminary results support the idea that the workflow performs well on other types of tissues, such as biopsies from solid tumors, effectively separating the relatively large numbers of cell types present in such tissues. I also did not present a benchmark the runtime of the proposed workflow as a function of dataset size. However, it is clear that the t-SNE step is much slower than any of the other steps, and the runtime of t-SNE on single-cell data has been benchmarked previously^13^. In our hands, t-SNE took approx. two minutes on the PBMC dataset with 10,392 cells. Our work highlights the ability to effectively analyze scRNA-Seq data by combining a simple normalization procedure with an appropriate variance-stabilizing transformation (VST) for Poisson-distributed data^4,11^. Future research focused on the development of effective methods for analyzing scRNA-Seq data could further explore the use of VSTs with the goal of designing simple and effective analysis workflows.

## Methods

### Description of the Galapagos workflow

Galapagos consists of a simple series of steps that takes as input a matrix containing UMI counts for *p* genes and *n* cells, and generates a two-dimensional t-SNE embedding of the data, as well as an assignment of cells in the dataset to clusters (a subset of cells may be designated “outliers” instead of being assigned to a particular cluster). I will first provide a description of each step, and then discuss the motivation and purpose of each step below. The Galapagos workflow consists of:

1. Median normalization. Let *C_i_* refer to the total UMI count of the *i*’th cell (*i*=1,…,*n*), and let *C* be the median across all total UMI counts: *C* = median(*C*_1_,…,*c_n_*). To normalize the expression profile of the *i*’th cell to *C*, the expression measurements (UMI counts) for all genes in that cell are then scaled by the factor (*C/c_i_*). The application of this scaling procedure to all cells results in a new dataset in which the total UMI count of the expression profiles of all cells equals *C*. This normalization procedure was proposed by Grun et al. in their analysis of noise models for scRNA-Seq data.
2. Variance stabilization. Let *X_ij_* refer to the normalized expression value of the *j*’th gene in the *i*’th cell. A new dataset with variance-stabilized values is then obtained by replacing the original value *X_ij_* with transformed values *y_ij_*, where 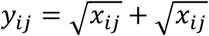. This transformation is known as the Freeman-Tukey transformation^9^ and effectively stabilizes the technical variance associated with scRNA-Seq measurements^4^.
3. Principal component analysis (PCA). Let **X** be the *n*-by-*p* matrix containing the normalized and variance-stabilized expression values. The first *d* principal components (PCs) are then determined by applying an efficient randomized algorithm for singular value decomposition (SVD)^15^ to **X**, producing a *p*-by-*d* matrix **W** whose columns are unit vectors that correspond to the first *d* principal components of **X**. The projection of **X** onto these PCs results in a new *n*-by-*d* matrix ***Y*** = ***XW*** that contains the PC scores of all cells for the first *d* PCs. The number of principal components *d* is determined by the user with a parameter called *num_components*. By default, Galapagos uses *num_components* = 30.
4. Visualization with t-distributed stochastic neighbor embedding (t-SNE)^7^. The Barnes-Hut implementation of t-SNE^16^ is applied to **Y**, resulting in an *n*-by-2 matrix **T** that contains the 2-dimensional coordinates of each cell in the t-SNE embedding. The *perplexity* parameter of t-SNE is determined by the user. By default Galapagos uses *perplexity* = 30.
5. Clustering with DBSCAN^10^. DBSCAN is applied to **T**, resulting in a variable number of clusters *k*, and a vector ***z*** = (*z_1_*,…, *Z_n_*) containing cluster assignments for each cell (*z_i_* ∈ [-1,1,2,…, k]), where a value of −1 indicates that a cell represents an outlier and does not belong to any cluster. DBSCAN has two parameters, *eps* and *minPts*, which have to be fine-tuned by the user in order to achieve results that are deemed consistent with the t-SNE visualization. As a starting point for the fine-tuning process, I suggest the use of *minPts* = [0.01 * nļ and *eps* = 0.03 * ‖*ν*‖, where ***v*** is a two-dimensional vector representing the range of x and y values (maximum - minimum) in the t-SNE visualization. In other words, *eps* is set to 3% of the “diameter” of the t-SNE plot.

The design of the Galapagos workflow was motivated by the following considerations:

1. The normalization step has two main purposes. First, it counteracts efficiency noise, i.e., technical fluctuations in mRNA detection efficiency between cells (droplets). Second, it ensures that the entire analysis is based on mRNA concentrations, instead on absolute mRNA counts. Cells of different types may exhibit differences in size and absolute mRNA content. As a result, gene concentrations (then number of mRNAs from “Gene X” relative to the total number of mRNAs detected), may be a better measure than absolute mRNA count for the effective availability of “Gene X” transcripts for translation into proteins.
2. The variance stabilization step relies on the observation that after normalization, the only major source of technical variability is sampling noise, which can be closely approximated with a Poisson model. It is known that for Poisson-distributed data, the Freeman-Tukey (FT) transform, 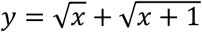, serves as an effective variance-stabilizing transformation (VST), at least for true values >= 1^9^. This ensures that the high absolute amounts of technical variance associated with highly expressed genes do not drown out the signal from more lowly expressed genes, essentially “leveling the playing field” for all genes. There is a concern that the expression differences of lowly expressed genes (e.g., with expression values < 0.1) remain significantly underrepresented, as differences in that range are highly compressed by the FT transform. However, due to the low detection rates of scRNA-Seq protocols, the signal-to-noise ratio for lowly expressed genes is generally very low. It therefore seems safe to assume that even with a less biased VST, these expression differences would not contribute greatly to a genome-wide analysis of expression differences.
3. After the data have been normalized and variance-stabilized, the projection of the data onto the first few (~30) PCs serves as an effective way to remove a large proportion of the technical noise present in the data. While cell type-specific expression patterns manifest in strong gene correlations, the technical noise does not exhibit any significant correlation structure. Therefore, the first few principal components can be expected to capture true expression differences between cells, while the higher principal components are expected to represent technical noise^11^.
4. t-SNE is a widely popular method for nonlinear dimensional reduction. The application of t-SNE to scRNA-Seq datasets that contain discrete cell types (as opposed to more continuous phenotypes) mainly serves to aggregate cells belonging to the same cell type, and to provide an overview of the number and size of cell populations contained in the data. t-SNE does not preserve global distances. Rather, it produces “cell islands” representing different cell types, and arranges those islands in a more or less arbitrary fashion within the two-dimensional plane. Importantly, t-SNE appears to produce islands with relatively uniform cell densities, which can lead to a “crowded” appearance when a large number of cells (e.g., 20,000 or more) is analyzed.
5. DBSCAN is a widely used density-based algorithm for clustering data. Abstractly, DBSCAN proceeds as follows: a) Construct a graph in which any two data points are connected by an edge if the distance between them is below a certain value (eps). b) Designate all data points with at least a minimum number of edges (minPts) as “core points”. c) Identify the connected components of all core points, ignoring all other points. Each connected component is a cluster. d) Assign all non-core points to the cluster closest to them, if the distance is below eps. e) Label all remaining points (if any) as outliers. DBSCAN performs well on t-SNE visualizations as the cell islands generally appear exhibit a uniform cell density, which often makes it possible to automatically assign the cells of each island with only one DBSCAN run. However, DBSCAN is limited by the extent to which t-SNE is able to clearly separate cells of different cell types. A more detailed discussion of the fine-tuning of DBSCAN parameters is provided below.

The tuning of the four parameters associated with the Galapagos workflow (*num_components, perplexity, eps*, and *minPts*) is straightforward. In the most situations, the default values of *num_components* (30) and *perplexity* (30) do not require adjustments. In cases where cell type heterogeneity is very low, it may be beneficial to lower the setting of *num_components* (e.g, to 10). Conversely, for tissues with a very large number of distinct cell types, it may be beneficial to increase *num_components* (e.g., to 50). As a rule of thumb, I suggest to allow for at least one PC per expected cell type (note that for very similar cell types, t-SNE may fail to effectively separate the cell populations, independently of how many additional PCs are included in the analysis). In general, including “too many” should not have a severe negative impact on the clustering results, since t-SNE is quite robust to the additional technical noise captured by the “superfluous” PCs. In contrast, including “too few” PCs can lead to the unintended combination of multiple cell types in one cluster, as the PCs capturing their differences have been excluded from the analysis. It is therefore advisable to always err on the side of including “too many” PCs. I have not yet encountered a situation where there was a clear rationale for changing the value of *perplexity*. It should be noted that increasing *perplexity* can lead to significantly longer t-SNE runtimes.

The tuning of the two DBSCAN parameters *eps* and *minPts* is rather intuitive in practice, as the DBSCAN algorithm is very fast, and the results can be easily judged by overlaying the cluster annotations onto the t-SNE result. The researcher can stop tuning whenever he/she is satisfied with the visual agreement between the cell islands in the t-SNE plot and the cluster assignments produced by DBSCAN. To simplify the testing of sensible parameter values, I suggest to reparametrize the algorithm by expressing *eps* as a fraction of the “diameter” of the t-SNE plot (with a default value of 3%), and *minPts* as a fraction of the total number of cells in the dataset (with a default value of 1%). These default values typically result in a relatively coarse clustering. For more granular clusters, *eps* can be decreased, which reduces the likelihood that core points belonging to different cell islands will become part of the same cluster (connected component). Whenever *eps* is reduced, it is advisable to also reduce *minPts*, to avoid a situation where most points fail to meet the criterion of a core point, and most cells are considered “outliers”. Conversely, when *eps* is increased, it might be useful to also increase *minPts*, to avoid an overly coarse clustering. Anticipating how different combinations of eps and *minPts* affect the DBSCAN clustering result can require some practice. To ensure that the results of each DBSCAN run can be examined quickly, I recommend to first obtain and store the t-SNE visualization result, so that different DBSCAN results can be visualized quickly without having to re-run the much more time-consuming t-SNE algorithm.

### Implementation of the Galapagos workflow

The Galapagos workflow was implemented in Python 3.7, using version 0.20.3 of the *scikit-learn* package^17^. PCA was performed using the *decomposition.PCA* class with *svd_solver*=‘randomized’. t-SNE was performed using the *manifold.TSNE* class. DBSCAN was performed using the *cluster.DBSCAN* class with *algorithm*=‘brute’. t-SNE and clustering results were plotted with plotly^18^ 3.7.0.

### Single-cell RNA-Seq preprocessing workflow

All scRNA-Seq datasets were preprocessed using a uniform pipeline, as previously described^11^. I reproduce the description of the procedure here for the reader’s convenience.

To ensure consistent preprocessing of all scRNA-Seq datasets analyzed in this study, I aimed to apply a uniform scRNA-Seq preprocessing workflow to all datasets. The goal of this workflow is to only retain known protein-coding genes that are located on the nuclear genome (not the mitochondrial genome), and to remove cells that do not meet certain quality control criteria. First, I extracted a list of known protein-coding genes from the human Ensembl genome annotations (e.g., release 92; http://ftp.ensembl.org/pub/release-92/gtf/homosapiens/Homosapiens.GRCh38.92.gtf.gz). I then curated a list of 13 protein-coding genes located on the mitochondrial genome as all genes that have a name starting with The first step of the preprocessing workflow consists of removing all genes that are not contained in the list of protein-coding genes (genes were identified using their Ensembl IDs). In a second step, I remove all cells with a total transcript count of less than 1,000 (counting only transcripts from the protein-coding genes). I also remove cells where the 13 mitochondrially encoded genes account for more than 15% of the observed transcripts. This removes cells with low-quality data, represented by a low total transcript count and/or a high proportion of mitochondrially encoded transcripts, potentially due to cell lysis.

### Demonstration of Galapagos on PBMC scRNA-Seq data

To demonstrate Galapagos on PBMC scRNA-Seq data (**Figure 1b,c**), I relied on a PBMC dataset labeled “10k PBMCs from a Healthy Donor (v3 chemistry)”, which belonged to a series of datasets made available by 10X Genomics under the heading “Chromium Demonstration (v3 Chemistry)” (https://www.10xgenomics.com/resources/datasets/). I downloaded the UMI-filtered expression matrix from the 10x Genomics website (http://cf.10xgenomics.com/samples/cell-exp/3.0.0/pbmc10kv3/pbmc10kv3filteredfeaturebcmatrix.tar.gz), and applied the scRNA-Seq preprocessing workflow (see above; using genome annotations from Ensembl release 97), which resulted in a dataset containing 10,392 with a median transcript count of 5831.0. I then applied Galapagos, with *eps*=5.49 (2. 6% of t-SNE diameter), *minPts=73* (0.7% of total cells), and default parameters otherwise.

The resulting clusters were manually annotated with the cell types shown in **Figure 1b** by examining the expression patterns of known expression markers and other genes with cluster-specific expression (**Figure 1c** and **Supp. Figure 1**). Two relatively small clusters exhibited expression of markers from multiple cell types and were therefore assumed to represent doublets.

### Evaluation of Galapagos on simulated scRNA-Seq PBMC data

To simulate scRNA-Seq data, I relied on a previously described approach^11^. I reproduce the description of this procedure in the following paragraph for the reader’s convenience.

To simulate realistic PBMC scRNA-Seq data, I decided to use real PBMC scRNA-Seq datasets as a template by applying ENHANCE^11^ and treating the resulting denoised matrix as the ground truth, and then simulating a noisy “raw” gene expression matrix with the same number of cells as the real dataset. To simulated efficiency noise, I first calculated a scale factor for each cell, by dividing the cell’s total transcript count in the real data by its total transcript count in the denoised data. I then obtained efficiency factors for each cell in the simulated dataset by applying bootstrapping (sampling with replacement) to this set of scale factors. I then scaled the expression profile of each cell in the ground truth by its simulated efficiency factor. After scaling the ground truth expression profiles to simulate efficiency noise, I obtained the simulated transcript count for the *j*’th gene in the *i*’th cell of the simulated matrix by sampling from a Poisson distribution with *λ=x_ij_*, where *x_ij_* represents the value of the *j*’th gene in the *i*’th ground truth expression profile (after scaling).

To define true cell type identities based on the ground truth, I applied Galapagos with eps=7.06 (2.7%) and minPts=42 (0.4%). To compare the total amounts of variance in the real, ground truth, and simulated datasets, I first median-normalized and FT-transformed each dataset, calculated the variance for each gene, and calculated the sum total (“true variance”).

### Evaluation of Galapagos on CITE-Seq PBMC data

To evaluate Galapagos on CITE-Seq PBMC data, I downloaded and preprocessed a PBMC CITE-Seq dataset labeled “10k PBMCs from a Healthy Donor - Gene Expression and Cell Surface Protein”, as described previously^11^. I reproduce the description of this procedure in the following paragraph for the reader’s convenience.

The dataset belonged to a series of datasets made available by 10X Genomics under the heading “Chromium Demonstration (v3 Chemistry)” (https://www.10xgenomics.com/resources/datasets/). I downloaded the UMI-filtered expression matrix and separated the transcriptome from the protein measurements, thus producing a transcriptome and a protein expression matrix (the measurements for the 17 proteins were easily identified as their gene identifiers were the only ones that did not start with “ENSG…”). I then applied the scRNA-Seq preprocessing workflow to the transcriptome expression matrix (see above; using genome annotations from Ensembl release 95), which resulted in a dataset containing 7,666 cells with a median transcript count of 3277.5. I then filtered the protein expression matrix to only retain the measurements from these cells.

To define T cells, NK cells, and monocytes based on the protein expression data (**Figure 4b**), I first normalized all protein expression profiles to 10,000 UMIs, to the protein expression matrix, and then defined T cells as cells with a CD3 count of at least 500, and a CD56 count between 3 and 3,000. This resulted in the identification of 4,111 T cells. Using the remaining cells, I then defined NK cells as cells with a CD56 count of at least 150, and a CD14 count of at most 200. This resulted in the identification of 954 NK cells. Using the remaining cells, I then defined monocytes as cells with a CD14 count of at least 1,000, or a CD16 count of at least 1,000. This resulted in the identification of 1,927 monocytes. This procedure left 674 cells without an identified cell type. This included B cells, for which no protein expression marker was available in the data. Since the identification of doublets based on protein expression data would have required individual gates for each type of doublet (e.g., T cell / CD14+ monocyte, T cell / CD16+ monocyte, T cell / NK cell, T cell / B cell), and as markers for B cells were not available, I instead opted to define as doublets the cells belonging to cluster 7 in the mRNA clustering result (**Supp. Figure 3a**), and to label those cells as “Other cells”, independently of their protein expression-based cell type assignments. The precision and recall values for each cell type (**Figure 4e**) were then calculated with the ‘precision_recall_fscore_support’ function from scikit-learn.

To define CD4+/CD8+ naïve T cells and CD4+/CD8+ memory T cells based on the protein expression data (**Figure 5a**), I again used the normalized expression profiles, and selected the previously defined 4,111 T cells. I then defined naïve T cells as cells with CD45RA count of at least 700, and a CD45RO count between 7 and 75. I also defined memory T cells as cells with a CD45RA count between 22 and 250, and a CD45RO count of at least 250. After selecting the naïve T cells, I defined CD4+ naïve T cells as cells with a CD4 count of at least 900, and a CD8a count between 10 and 95. I also defined CD8+ naïve T cells as cells with a CD4 count between 6 and 90, and a CD8a count of at least 1,000. After selecting the memory T cells, I defined CD4+ memory T cells as cells with a CD4 count of at least 900, and a CD8a count of at most 95. I also defined CD8+ memory T cells as cells with a CD4 count of at most 90, and a CD8a count of at least 1,000.

I applied Galapagos to the mRNA expression data (**Figure 4a,d**) with *eps*=5.87 (2.7%) and *minPts=68* (0.09%), and default parameter values otherwise. To apply Galapagos on only the T cells (**Figure 5c**), I first selected the 4,111 T cells, and again excluded the cells defined as doublets (see above). I then obtained a t-SNE visualization (**Figure 5b**) by applying Galapagos with default parameters.

To visualize the mRNA expression data with UMAP (**Supp. Figure 5**), I replaced t-SNE with UMAP in the Galapagos workflow, and again used the default values of 30 for *num_components*. I applied UMAP using the Python package umap-learn 0.3.8, with the parameters *num_neighbors=15* (the default value) and *min_dist=0.5* (default value: 0.1) and default values otherwise.

## Acknowledgements

I drew inspiration for this work from a presentation that I was invited to give at the Applied Bioinformatics Lab’s Single-cell RNA-Seq Analysis Club at NYU School of Medicine in February 2019. I would therefore like to thank the organizers and participants of the event. In particular, I would like to thank my advisor Itai Yanai for the invitation.

## Notes

### Competing Interest Statement

The authors have declared no competing interest.

### Summary of Updates

Updated Discussion section.

