## Supplemental Figures for "Straightforward clustering of single-cell RNA-Seq data with t-SNE and DBSCAN"

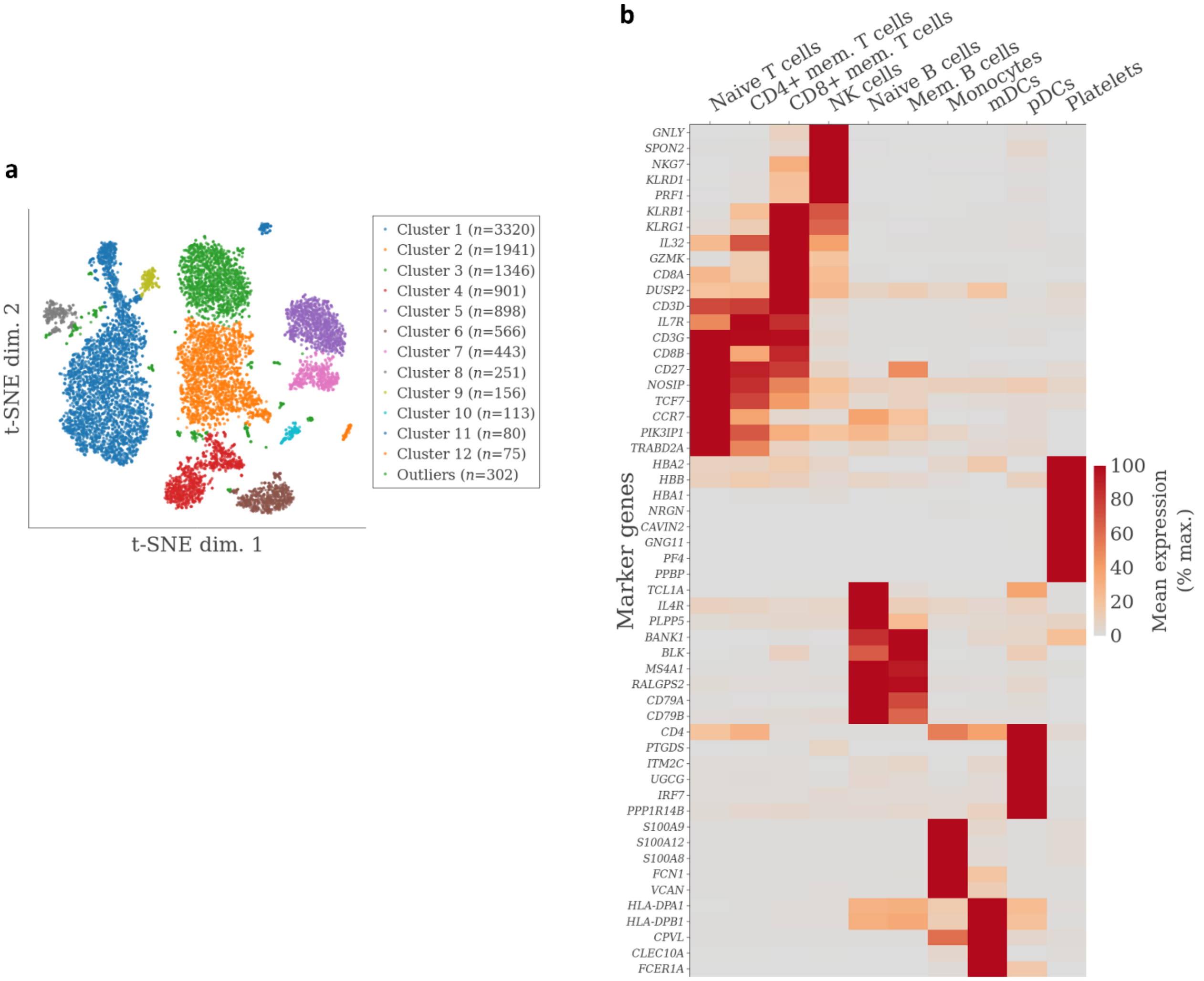


**Supplementary Figure 1: Galapagos results for scRNA-Seq PBMC data. a** Original Galapagos result that formed the basis for the result shown in **Figure 1b** (right). **b** Expanded list of genes with cell type-specific expression patterns. See **Figure 1c** for legend.

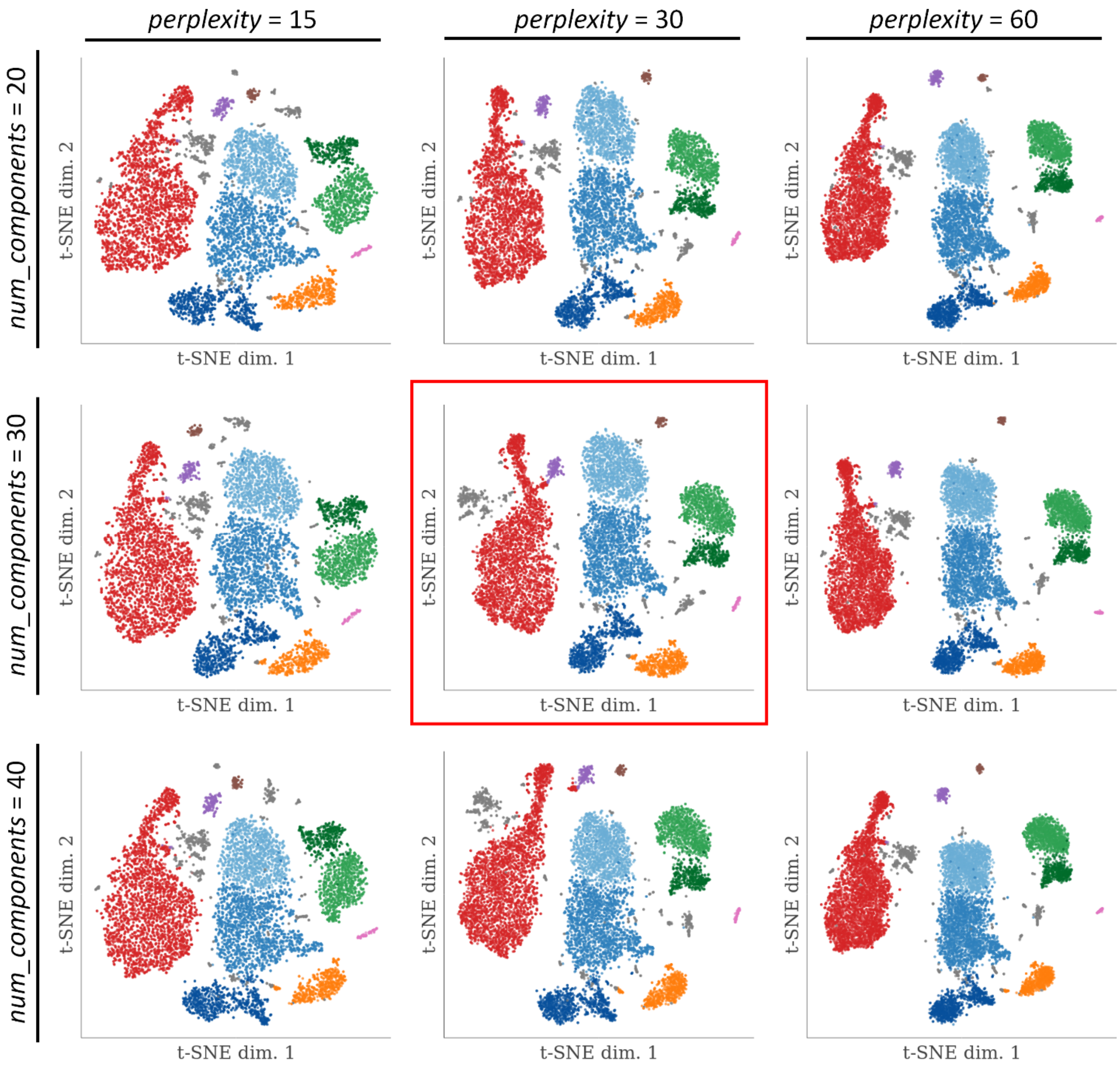


**Supplementary Figure 2: Robustness of PBMC t-SNE results to parameter choices.** The values for *num_components* and *perplexity* are indicated, and the result in the center (red box) is identical to that shown in **Figure 1b** (right). In all other plots, the cluster assignments obtained in this analysis were overlaid.

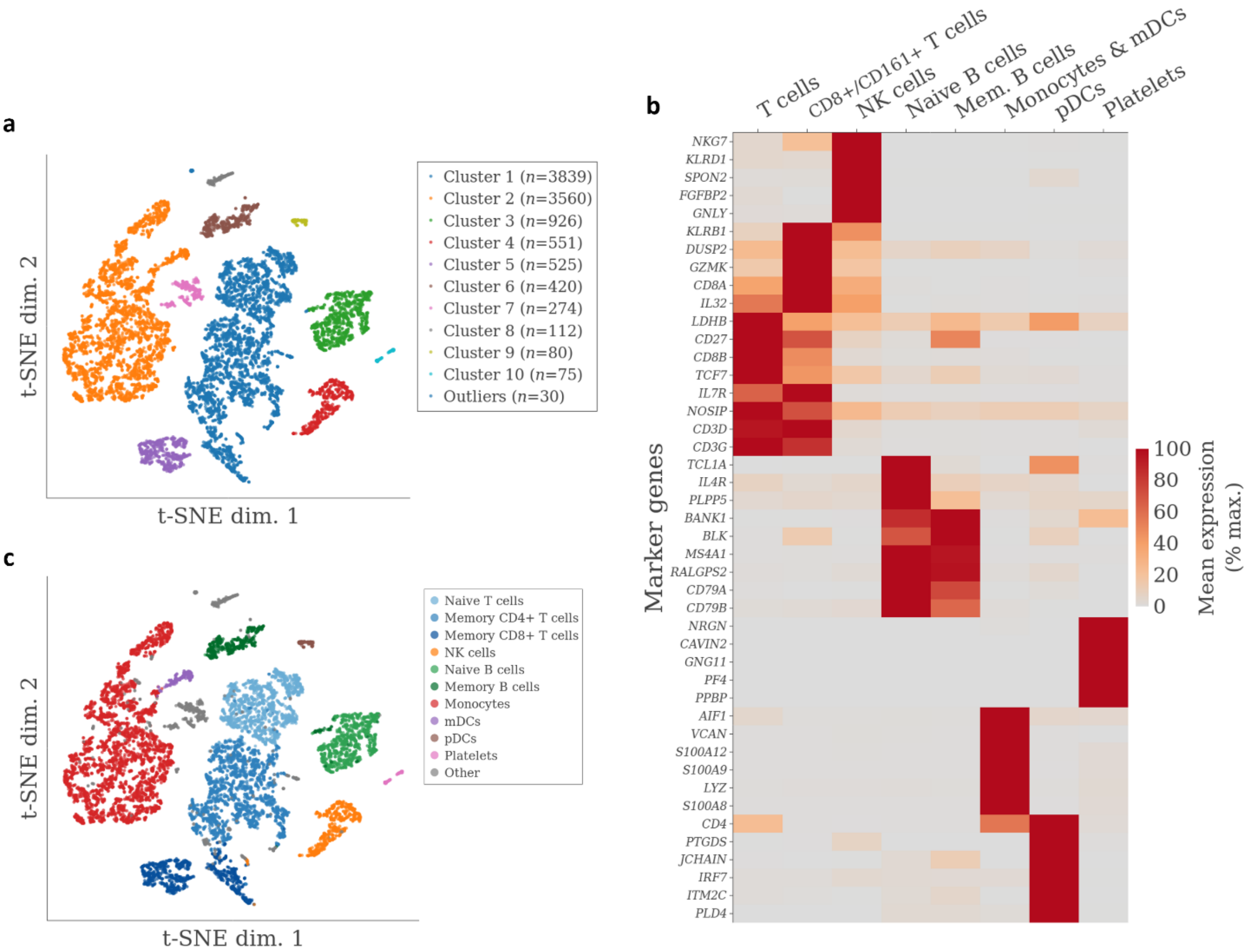


**Supplementary Figure 3: Galapagos results for simulated scRNA-Seq PBMC data. a** Original Galapagos result for the ground truth data. This result formed the basis for the result shown in **Figure 3b**. **b** Expanded list of genes with cell type-specific expression patterns. See **Figure 1c** for legend. **c** Same t-SNE visualization as in (**a**), but overlaid with the clustering results obtained for the real dataset (**Figure 1b**, right). This is possible since there is a 1:1-correspondance of cells between the real and the simulated data.

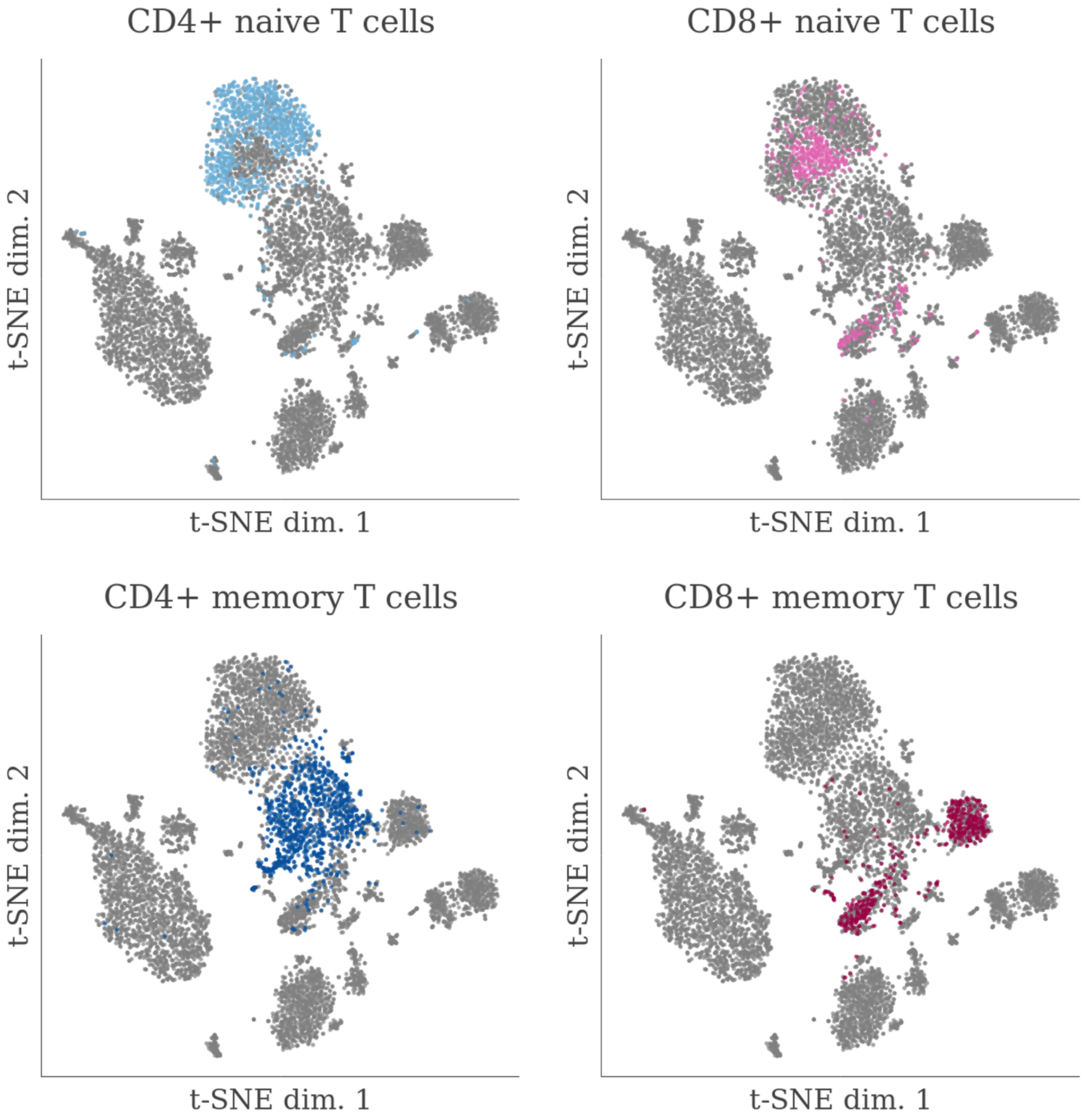


**Supplementary Figure 4: Detailed view of the agreement between t-SNE results and CITE-Seq-derived T cell subtype identities.** Shown are the same results as in **Figure 5b**, but the cells assigned to each cell type are highlighted in a separate plot for improved readability.

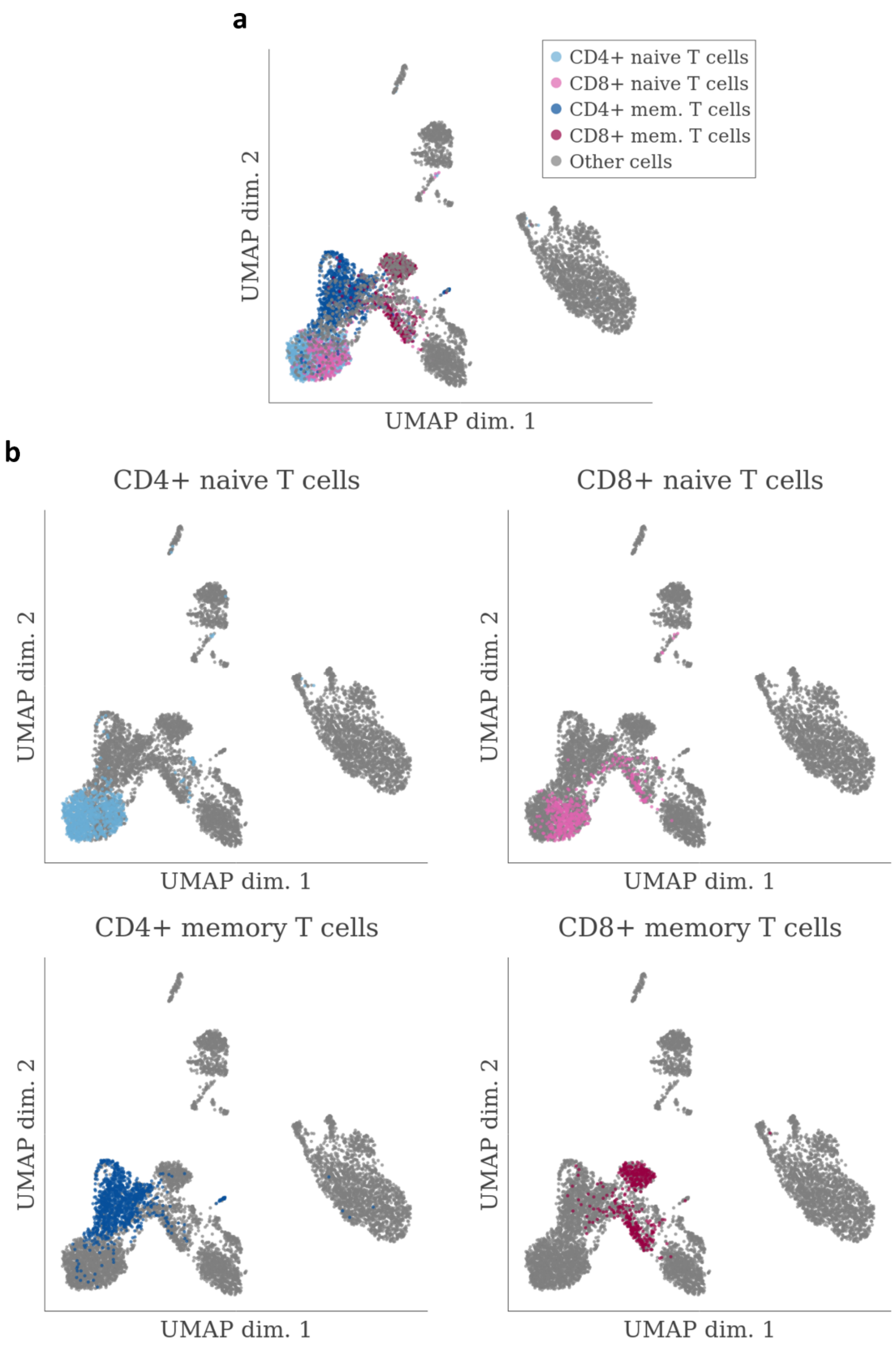


**Supplementary Figure 5: Application of UMAP to PBMC data and comparison with CITE-Seq derived T cell subtype identities. a** UMAP visualization overlaid with CITE-Seq derived T cell subtype identities. **b** Detailed views where only cells annotated with one T cell subtype are highlighted for improved readability.
